# Zebrafish model of human Zellweger syndrome reveals organ specific accumulation of distinct fatty acid species and widespread gene expression changes

**DOI:** 10.1101/2021.01.03.425169

**Authors:** Shigeo Takashima, Shoko Takemoto, Kayoko Toyoshi, Akiko Ohba, Nobuyuki Shimozawa

## Abstract

In Zellweger syndrome (ZS), lack of peroxisome function causes physiological and developmental abnormalities in many organs such as the brain, liver, muscles, and kidneys, but little is known about the exact pathogenic mechanism. By disrupting the zebrafish *pex2* gene, we established a disease model for ZS and found that it exhibits a pathological condition and metabolic failures similar to that of human patients. By comprehensive analysis of fatty acid profile, we found organ specific accumulation and reduction of distinct fatty acid species such as an accumulation of ultra-very-long-chain polyunsturated fatty acids (ultra-VLCPUFAs) in the brain of *pex2* mutant fish. Transcriptome analysis using microarray also revealed mutant-specific gene expression changes that might lead to the symptom, which include reduction of *crystallin, troponin, parvalbumin*, and fatty acid metabolic genes. Our data indicated that the loss of peroxisome results in widespread metabolic and gene expression changes beyond the causative peroxisomal function. These results suggest the genetic and metabolic basis of the pathology of this devastating human disease.

## INTRODUCTION

Peroxisomes are small vesicular eukaryotic organelles that are found in the cytoplasm, and are metabolic foci of many biological molecules. Examples of peroxisomal metabolic functions include degradation of very-long-chain fatty acids (VLCFAs: fatty acids with 22 or more carbon atoms), degradation of branched fatty acids such as phytanic acid and pristanic acid, the first steps of ether phospholipid synthesis (including plasmalogens), bile acid synthesis, and hydrogen peroxide decomposition. In primates, docosahexaenoic acid (DHA) synthesis also requires peroxisomes (Morita and Imanaka, 2020). Peroxisomal metabolism is essential for human health and its disorders lead to various serious health problems.

Peroxisome dysfunction due to genetic alterations underlies congenital metabolic diseases collectively called peroxisomal disorders (Shimozawa, 2020a; Wanders et al., 2016). Peroxisomal disorders comprise two major categories: peroxisome biogenesis disorder (PBD), which is associated with a failure of the cells to biosynthesize peroxisomes, broadly curtailing peroxisomal function, and single peroxisomal enzyme deficiencies, which reduce or eliminate a single or limited peroxisomal metabolic function(s). Peroxins, or peroxisomal biogenesis factors (PEXs) coordinate peroxisomal biosynthesis. Twelve known loci (*PEX1, 2, 3, 5, 6, 10, 12, 13, 14, 16, 19*, and 26) are associated with causative mutations underlying Zellweger spectrum disorder (ZSD), which is a severe subgroup of PBD. In ZSD, peroxisomal biosynthesis is strongly affected, disrupting all peroxisome dependent metabolic functions. Two loci (*PEX7* and *PEX5 long isoforms*) are known to be responsible for another subgroup of PBD, rhizomelic chondrodysplasia punctata (RCDP) types 1 and 5, in which selected metabolic pathways are affected. Sixteen loci are associated with peroxisomal single-enzyme deficiencies, each of which affects one to a few peroxisomal metabolic pathway(s) (Shimozawa, 2020a; Wanders et al., 2016).

Defective peroxisomes affect fetal and neonatal development in humans (Klouwer et al., 2016; Shimozawa, 2020a; Steinberg et al., 2006). Patients with PBD show multiple symptoms, including craniofacial dysmorphism, abnormal cerebral and cerebellar development with neuronal migration defects and irregular gyri. These patients also show liver defects, such as hepatomegaly and jaundice, renal cysts, glaucoma, retinitis pigmentosa, severe hypotonia, chondrodysplasia punctate, psychomotor retardation, and sucking disability. Three major forms of ZSD, Zellweger syndrome (ZS), neonatal adrenoleukodystrophy (NALD), and infantile Refsum disease (IRD) comprise a disease spectrum in which ZS is the most severe, NALD is less severe, and IRD is the least severe. The severity of disease correlates with the degree of gene disruption. The biochemical consequences of faulty peroxisomal metabolism include accumulation of VLCFAs, phytanic acid, and pristanic acid, depletion of ether phospholipids (plasmalogens) and bile acids. The range of metabolic fluctuation also correlates with the degree of gene disruption; however, the direct relationship between affected metabolites and disease pathogenesis is still unclear, hindering detailed understanding of this disease, and development of effective clinical treatments.

To address this problem, a disease model is required. To date, the effects of eliminating peroxisomes have been studied in animal models including mice, zebrafish, fruit flies, and nematodes (Baes and Van Veldhoven, 2012; Takashima and Shimozawa, 2020). In particular, several lines of *Pex* gene knockout mice have been produced (Baes et al., 1997; Faust and Hatten, 1997; Hanson et al., 2014; Hiebler et al., 2014; Li et al., 2002; Maxwell et al., 2003). Mutant mice show similar metabolic changes to those seen in human patients, such as increased VLCFAs, phytanic acid, and pristanic acid, and reduced plasmalogen and ether phospholipid levels. Mutant mice also recapitulate some symptoms of ZS, such as neuronal migration defects in the cerebral cortex, abnormal cerebellar development, severe hypotonia, lack of sucking ability, and liver abnormality, indicating a conserved role of peroxisomes in embryonic and postnatal development among mammalian species.

In zebrafish, Krysko and colleagues reported a Morpholino oligo-mediated *pex* gene knockdown (Krysko et al., 2010), where the peroxisome biogenesis was only partially disturbed probably because of remaining *pex* gene activity. Therefore, we preferred a gene knockout approach to eliminate peroxisome biogenesis. Here, we report the first gene-knockout model of ZS in zebrafish. We targeted the zebrafish *pex2* gene, one of the *pex* genes responsible for ZS, to eliminate peroxisome biogenesis during development. We found that the ZS model fish successfully eliminated peroxisomes, and displayed phenotypes similar to human patients with ZS. Our findings confirm the conserved role of peroxisomes in the development among lower to higher vertebrate species. We also performed comprehensive fatty acid analysis using liquid chromatography-mass spectrometry (LC-MS) and whole transcriptome analysis by gene expression microarray to search for metabolites and gene expression changes related to pathogenesis.

## RESULTS

### Generation of *pex2*-deficient ZS model fish by genome editing

To establish a Zellweger syndrome (ZS) disease model, we introduced a mutation in the zebrafish *pex2* gene, one of the disease-causing *PEX* genes in humans (Shimozawa et al., 1992). Human PEX2 protein is an E3-ubiquitin ligase with a zinc RING finger domain that resides in the peroxisomal membrane (Platta et al., 2009). A complex of PEX2, PEX10, and PEX12 proteins recycles PEX5 protein, which is required for translocation of peroxisomal matrix proteins and enzymes from the cytoplasm into the peroxisomal matrix (Kawaguchi and Imanaka, 2020). In the absence of PEX2 function, the import of peroxisomal matrix proteins is severely impaired, resulting in formation of defective peroxisomes devoid of matrix proteins (so-called ‘ghost peroxisomes’) that lack metabolic activities. In the zebrafish genome, there is one human *PEX2* orthologue. The sequence similarity between human PEX2 and zebrafish Pex2 proteins is 57% at the amino acid level. Both human and zebrafish *PEX2* genes are composed of two exons. While human PEX2 protein is translated exclusively from exon 2, zebrafish Pex2 is translated starting near the 3’ terminus of exon 1, from which only 4 nucleic acids contribute to the protein. We designed a pair of TALENs (TAL effector nucleases) to induce a double strand break at the anterior region of exon 2 of the zebrafish *pex2* gene (Fig. 1A; the target sequences of TALENs are shown in Supplementary Table S1). Synthetic mRNAs encoding the TALENs were microinjected into wild-type zebrafish embryos (RIKEN strain) at the 1-cell stage. The injected G0 embryos were reared to adulthood, and F1 progeny were recovered by cross mating the G0 fish. The *pex2* genomic fragment was amplified by PCR from genome DNA of F1 progeny, and DNA sequences were examined to determine whether they possessed in-del mutations. Founder F1 individuals were crossed with wild-type fish to expand the population. We have established two mutant lines, *pex2*^*typeB*^ and *pex2*^*typeC*^, each of which possesses distinct frame-shift mutations. In each frameshift mutation, a *de novo* stop codon appears shortly downstream of the mutation site, resulting in premature termination of translation (Fig. 1B). Because they are both considered null alleles, we primarily used *pex2*^*typeC*^ in the following studies, unless otherwise stated.

**Fig 1.**
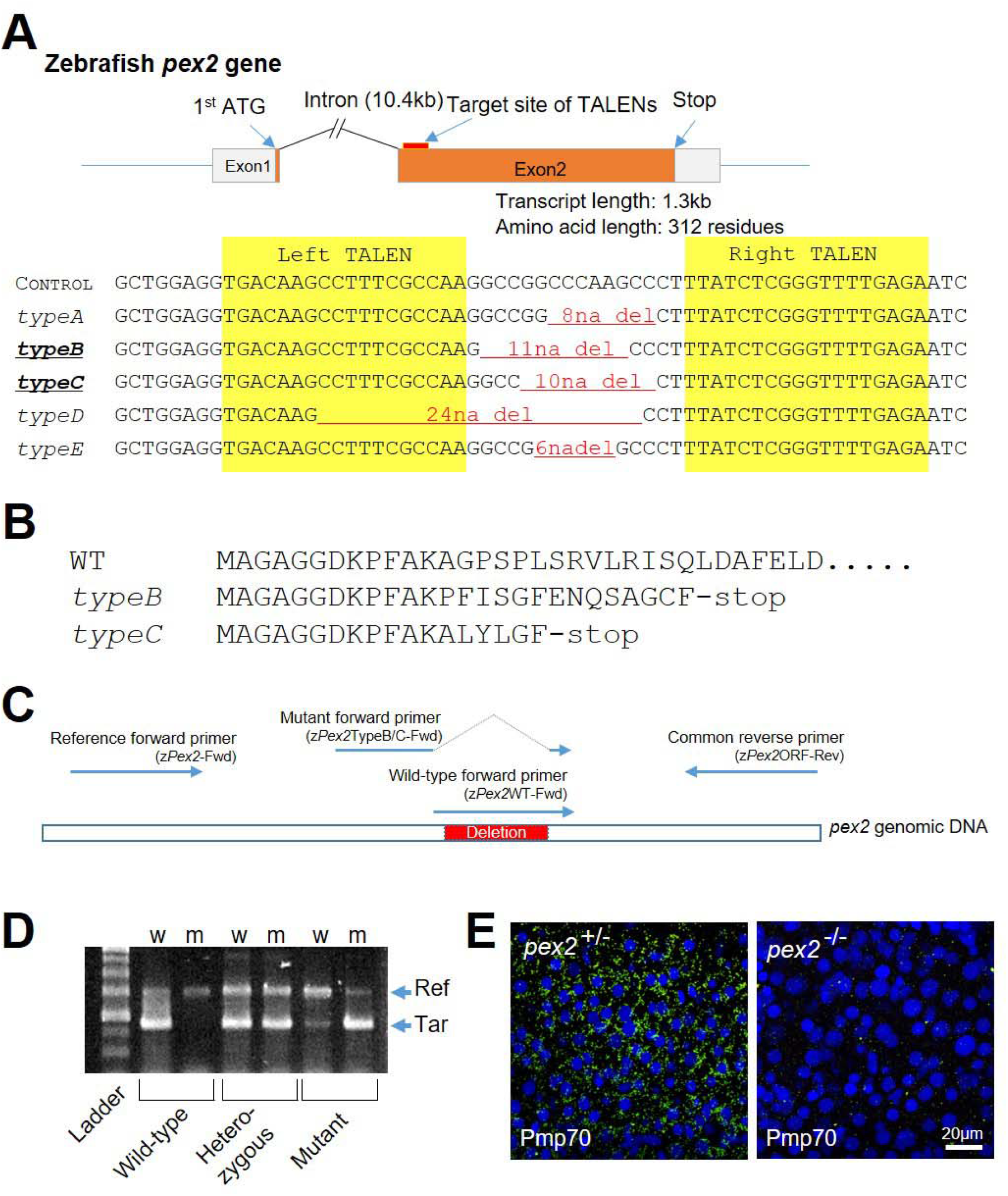
Generation of ZS model fish by TALEN-mediated mutagenesis. (A) TALEN Target sequences on the zebrafish *pex2* gene, and induced deletions. The zebrafish *pex2* gene contains two exons with the translation initiation site at the 3’ terminus of exon 1. Five deletions obtained at the F1 stage by TALEN-mediated mutagenesis are shown. We retained *typeB* and *typeC* alleles for this study. (B) Comparisons of predicted amino-acid sequences of mutated *pex2* genes. (C, D) Genotyping of mutant fish by genomic PCR. (C) Distribution and the name of the primers on the *pex2* locus is shown. (D) The genotypes of individual fish were determined by genomic PCR using wild-type (w) or mutant (m) specific primers and visualized by gel electrophoresis. The genotype was determined by the amplification pattern of the target fragment (Tar) and reference fragment (Ref). (E) Peroxisomal phenotype in the liver of a *pex2* mutant fish (right) and a wild-type sibling (left). Livers were stained with anti-Pmp70 (green) antibody. Cell nuclei were stained with DAPI (blue).

### Mutant fish show loss of peroxisomal structure, and high mortality

We confirmed the genotype of the mutant fish by genomic PCR. Primers were designed to amplify DNA fragments from the *pex2* locus. We designed two types of forward primer (a wild-type-specific primer and a mutant-specific primer) to distinguish wild-type and mutant *pex2* alleles (Fig. 1C and Table S1). Because mutant fish harbor 11 (*typeB*) or 10 (*typeC*) nucleotide deletions, the wild-type forward primer that includes the deleted nucleotides can only amplify genomic fragments from the wild-type allele. Mutant forward primers (*typeB* allele-specific and *typeC* allele-specific), which incorporate nucleotides flanking the deletions, were used to amplify mutant alleles (Fig. 1C). Using these primers, we were able to distinguish the genotype of the fish (Fig.1D).

Mutant embryos and larvae did not show clear external morphological abnormalities; however, we realized that most of the mutant fish died during growth. We examined the survival rate of the mutant fish and found it to be 50% by week two post fertilization, and 14% by the end of the second month (Table 1). Mutant fish rarely survived for more than two months. By microscopic examination, dying mutant larvae were found to contain little or no food in their digestive tracts, showing signs of an eating disability, which is often seen in human ZS patients. We examined livers of mutant and wild-type fish at three weeks by staining peroxisomes with antibodies against Catalase and Peroxisomal membrane protein 70 (Pmp70; also known as ATP-binding cassette sub-family D member 3, Abcd3). Catalase is a peroxisomal matrix protein that degrades hydrogen peroxide generated in peroxisomes. Pmp70 is a membrane protein that transports unsaturated fatty acids into peroxisomes for degradation by β-oxidation. Without PEX2 activity, import of peroxisomal matrix proteins is perturbed, but translocation of peroxisomal membrane proteins remains unaffected. In cultured fibroblasts obtained from ZS patients, matrix proteins such as catalase stay in the cytoplasm, while membrane proteins such as PMP70 reside in the membranes of the ghost peroxisomes. In wild-type (*pex2* ^+/+^ or *pex2* ^+/-^) livers, Catalase (data not shown) and Pmp70 (Fig. 1E, left panel) were localized to peroxisomes. In *pex2* ^-/-^ livers, unexpectedly, neither Catalase nor Pmp70 were detectable, showing complete loss of peroxisomal structure, including ghost peroxisomes, in contrast to the observation in human fibroblasts (Fig. 1E, right panel). Extensive loss of peroxisomal structure (including ghost peroxisomes) in the liver has been reported in human patients and ZS mouse models (Baes et al., 1997; Roels et al., 1995). Peroxisomes are entirely lost in the livers of *pex2* mutant fish, revealing successful elimination of peroxisomal biogenesis in *pex2* mutant fish.

**Table 1.**
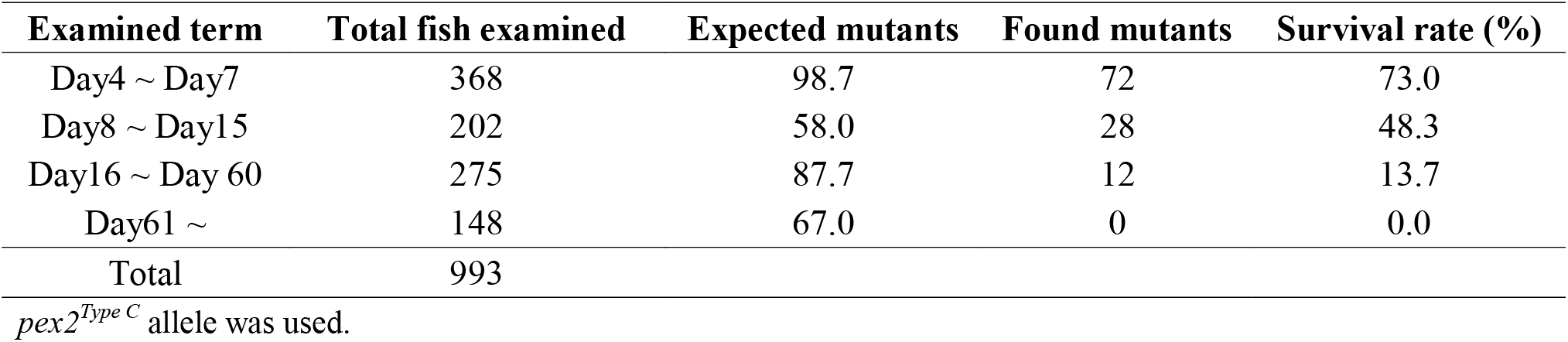
Survival rate of *pex2* mutant fish.

### Identification of FA species with organ-specific altered levels in mutants

In human ZS, changes in peroxisomal metabolite levels have been found. To elucidate organ-specific FA fluctuations in mutant fish, we compared FA profiles of brain, liver, and eyes between mutant and wild-type fish, and found that distinct FA species accumulated or reduced in each organ. Total FAs were extracted from these organs and analyzed by LC-MS. FA species were distinguished according to the carbon chain length and the number of double bonds. The amount of each FA species was shown as its ratio to total FAs detected (Fig. 2A). In control livers, we detected 49 FA species, ranging from C14 to C36. In mutant livers, 56 FA species were found. Compared to the control, mutant livers accumulated saturated VLCFAs as well as mono- and di-unsaturated VLCFAs (i.e., chain lengths of 22-36, with 0-2 double bonds). In the eyes, 86 and 100 FA species were found in control and mutant fish, respectively. In the brain, 89 and 111 FA species were found in control and mutant fish, respectively. In contrast to the liver, accumulation of highly polyunsaturated VLCFAs with chain lengths of 26-36 and 4-9 double bonds was prominent in mutant brains and eyes. In contrast, some of the shorter VLCFAs with no or few double bonds (C24-28, 0-3 double bonds) were decreased in the brain of mutant fish. These data reveal that the FA profiles differ between organs, and uncommon VLCFA species with much longer chain lengths (C32+) and higher double bond numbers (7+ double bonds), which we call ultra-very-long-chain polyunsaturated FAs (ultra-VLCPUFAs) (Takashima et al., 2017), accumulate in mutant organs.

**Fig 2.**
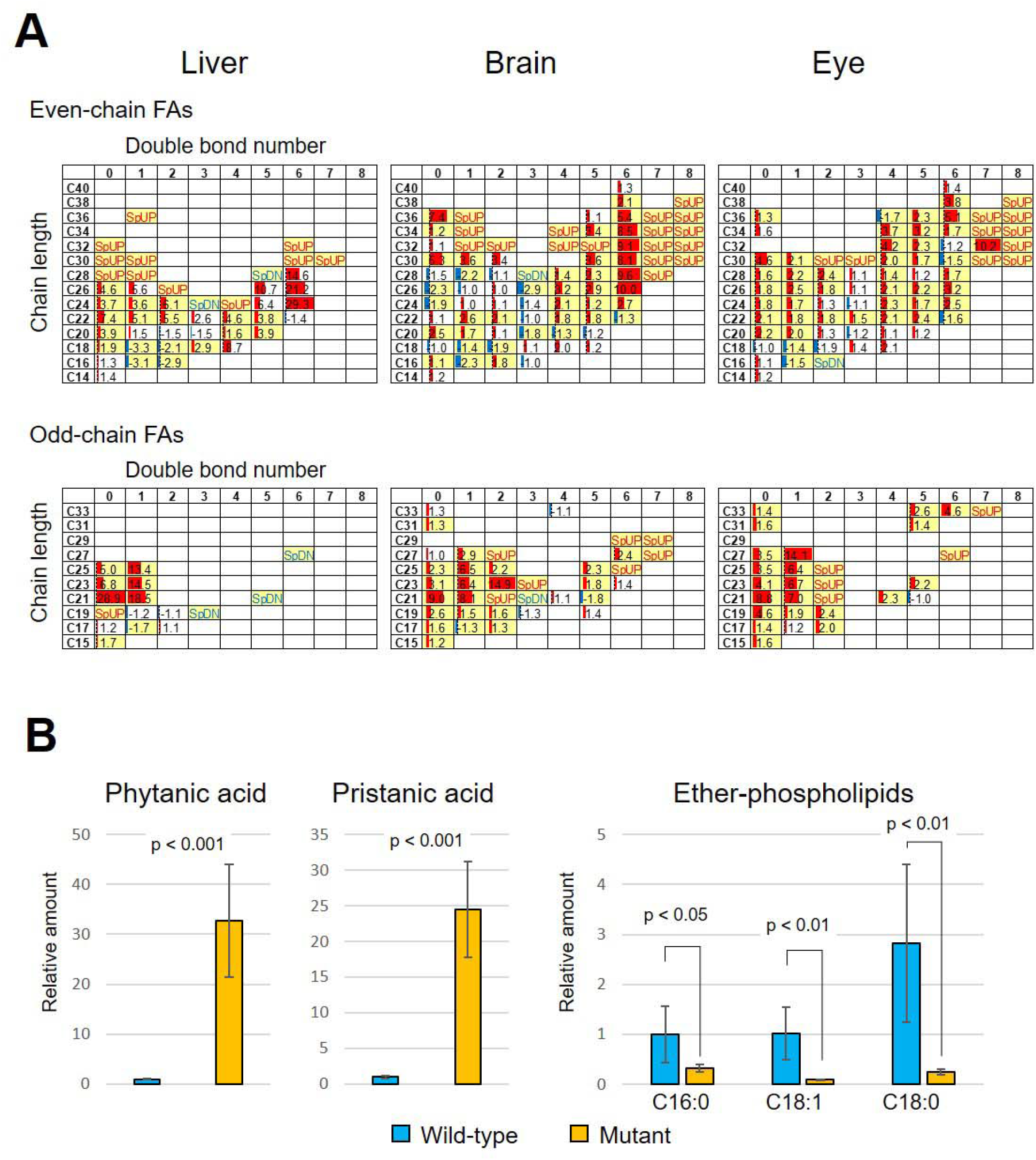
Analysis of FA profiles of mutant fish. (A) Comparison of FA profiles in liver, brain, and eyes between *pex2* mutant fish and wild-type siblings (n = 5 each). FA species were distinguished by chain length and number of carbon-carbon double bonds. Amount of FA species was compared between *pex2* mutant and wild-type siblings, and shown in color-coded data bars according to their abundance in the mutant fish. The number in the cells indicate the fold-change value. Student’s *t*-test and Benjamini-Hochberg multiple testing correction were applied to reveal significant quantitative differences between the two groups. An FDR cut-off of 0.05 was applied. The cells with yellow background indicates FA species with statistically significant increases or decreases. The cells labeled “SpUP” indicate FA species that were specifically detected in mutants, while the cells with “SpDN” indicate FA species that were specifically detected in wild-type fish. (B) Comparisons of branched FAs and ether phospholipids. The amount of phytanic acid and pristanic acid in the mutant liver were increased more than 30-fold or 20-fold, respectively. Ether phospholipids were detected in the form of 1-alkyl-sn-glycerol 3-phosphates possessing alkyl chain of C16:0, C18:1, and C18:0 fatty alcohol. The amount of ether phospholipids in the brain was significantly reduced in mutant.

We also found that branched fatty acids, such as phytanic acid and pristanic acid were highly accumulated in mutant liver, and ether phospholipids, detected in the form of 1-alkyl-sn-glycerol 3-phosphates (Takashima et al., 2017), were greatly reduced in mutant brain (Fig. 2B). Together, the data confirmed that the metabolic functions of peroxisomes are largely impaired in mutant fish.

### Mutant fish develop liver steatosis

Human ZS patients and model mice manifest liver defects, which we found in mutant fish as well. In human patients, liver biopsies have revealed fibrosis and cirrhosis. In ZS mice, hepatic steatosis and hepatitis are reported (Baes and Van Veldhoven, 2016). In normal zebrafish development, the primordial liver emerges around 4 days post fertilization (dpf), and continues to grow as the fish further develop. After 5 dpf, the liver primordium is clearly visible under a dissection microscope as semi-transparent tissue. We found some embryos showed decreased transparency (cloudy liver; Fig. 3A). Genotyping revealed that most (90%) of the *pex2*^-/-^ mutant embryos showed this cloudy liver phenotype (Fig. 3B). We stained embryos with Oil Red O (ORO), which specifically stains neutral lipids, and found strong staining in mutant livers (Fig. 3C). Using confocal microscopy, we observed large ORO stained droplets containing neutral lipids inside mutant hepatocytes. Droplets were much smaller, if any, in wild-type hepatocytes. Lipid droplets became larger as surviving mutant fish grew, and after 1 month, droplets reached a size as large as or larger than cell nuclei (Fig. 3D). In adults, the livers of normal fish have a three lobed appearance, with a pinkish color. In mutant fish, while the size was unchanged (Supplementary Fig. S1), the livers were highly abnormal in shape and deformed, lacked clear lobation, and were whitish in color, indicative of tissue inflammation (Fig. 5A). Accumulation of lipid droplets, a sign of liver steatosis, is also found in ZS model mice (Dirkx et al., 2005; Keane et al., 2007; Maxwell et al., 2003). Therefore, hepatic lipidosis must be a common phenotype of defective peroxisomal biogenesis among vertebrate species.

**Fig 3.**
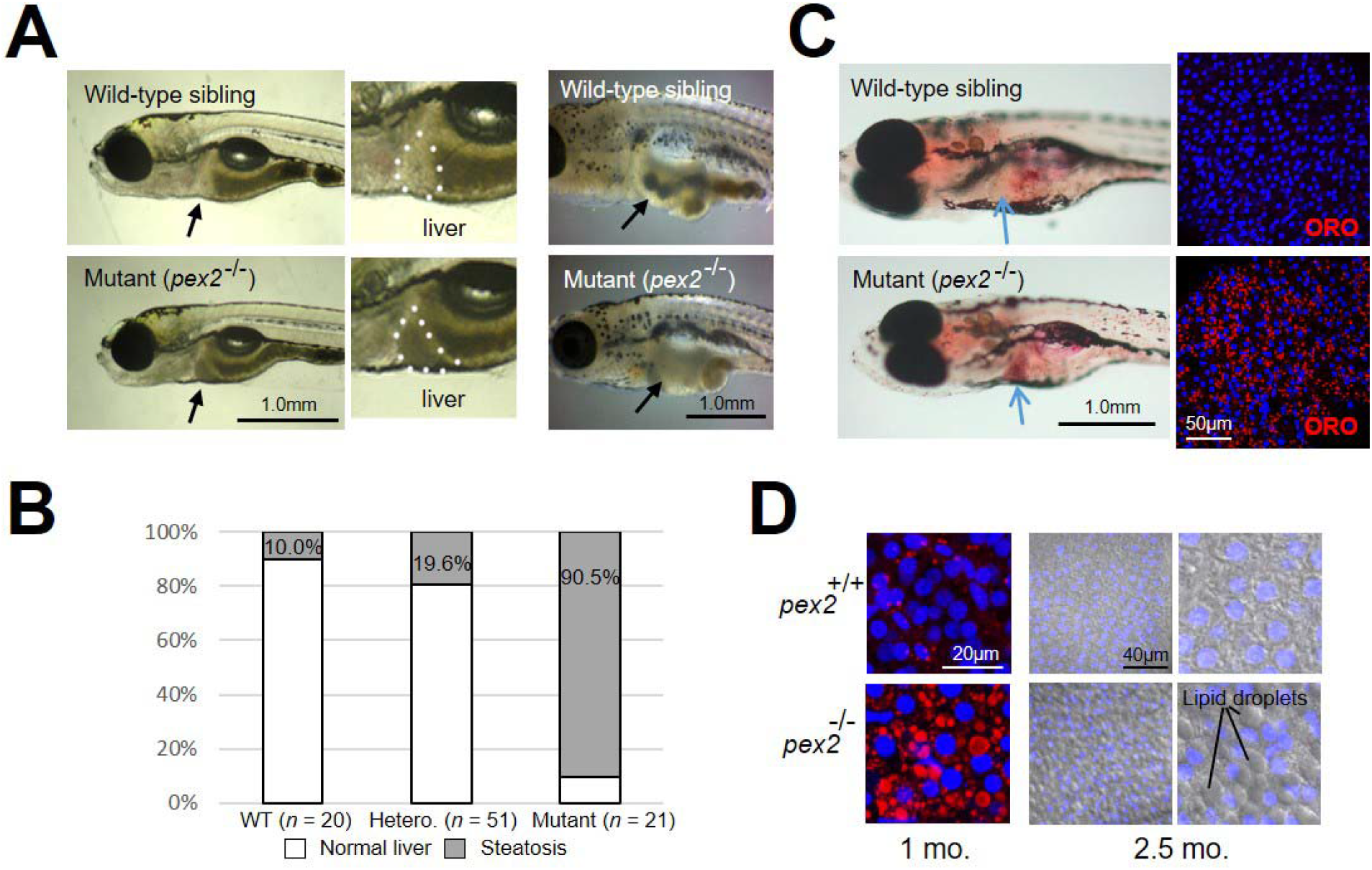
Liver phenotype of *pex2* mutant fish. (A) Livers of *pex2* mutant fish showed decreased transparency, which is a sign of steatosis. Larvae at 5 dpf (left four panels) and young adult fish at 1.5 months old (right two panels; the skin was partially removed) are shown. Livers are indicated by arrows or dotted lines. In the right pannels, intestine is visible through the semi-transparent liver in wild-type fish, while invisible in mutant. (B) The frequency of liver steatosis in *pex2* mutant fish is about 90% at 5-6 dpf. In wild-type (WT) and heterozygous fish (Hetero), it is below 20%. (C) Whole larvae at 14 dpf were stained with Oil Red O (ORO) reagent. Arrows indicate the position of the liver. Right panels show confocal microscopic images of dissected livers. Cell nuclei are in blue (DAPI staining). (D) Livers of older fish show large lipid droplets in their hepatocytes. Livers were stained with ORO and DAPI (1 month old, right panels) or DAPI alone (2.5 months, left four panels; lipid droplets are visible by transmitting light as large elliptical spheres).

**Fig 4.**
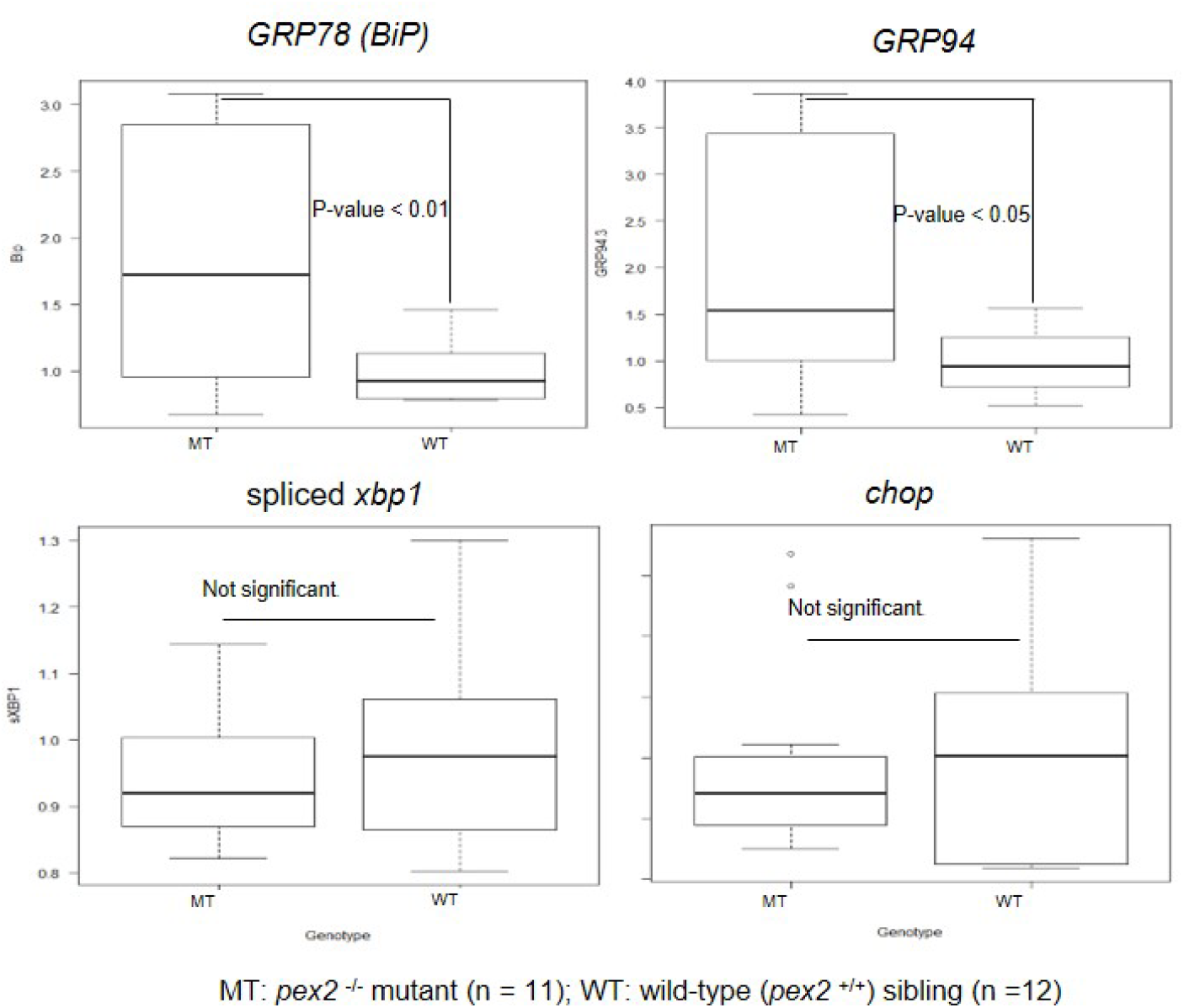
Real-time qPCR of ER stress-related genes. The relative amounts of mRNAs of ER-residing chaperones (*grp78*/*BiP* and *grp94*), the spliced form of *xbp1*, and *chop* were compared between *pex2* mutant fish (MT) and wild-type siblings (WT). Chaperone expression significantly increased in *pex2* mutant fish.

**Fig 5.**
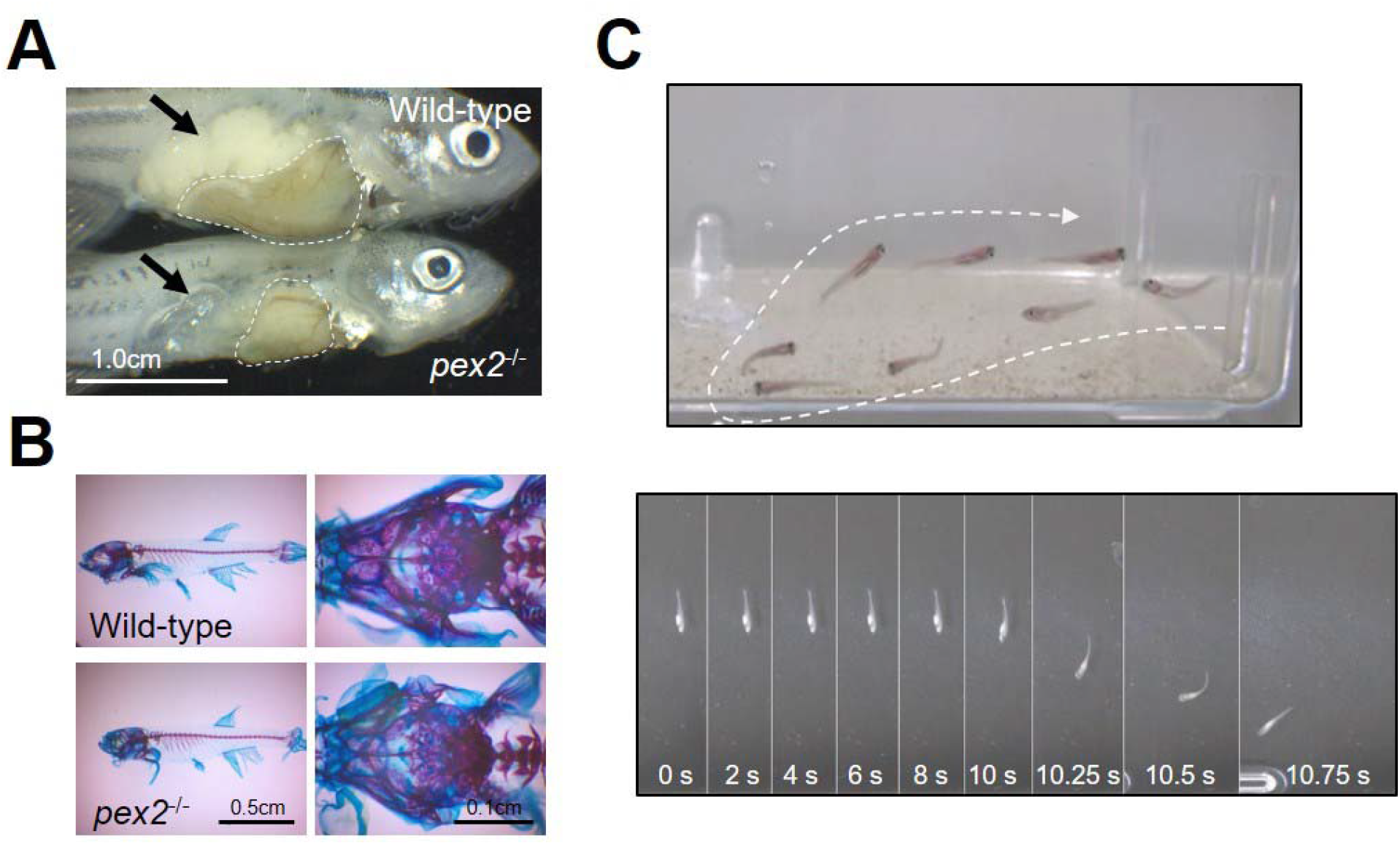
Internal phenotype and behavioral abnormality of adult mutant fish. (A) Two month-old fish were fixed and partially dissected to reveal internal organs. Livers are indicated by dotted lines, and ovaries are indicated by arrows. Note that livers of *pex2* mutant fish are deformed and whitish in color. The ovaries of *pex2* mutants lack mature oocytes, while those of wild-type siblings are full of them. (B) Skeletal specimen of adult fish stained by alizarin red S (bones) and Alcian blue (cartilages) revealed no skeletal abnormality in mutant fish. Skulls were magnified in right panels. (C) Abnormal behavior of mutant fish. (Top) Superimposed image of mutant fish showing swimming difficulty. Note the poor body control while swimming. Arrows indicate the moving direction. (Bottom) Time-lapse image of a mutant fish. The numbers indicate the elapsed time. The fish has lain on the bottom of a fish tank and started moving when the tank was tapped at 10s.

### Elevated ER stress and increased ER chaperones in mutant livers

Liver steatosis correlates with elevated stress in the endoplasmic reticulum (ER) (Han and Kaufman, 2016; Lebeaupin et al., 2018; Malhi and Kaufman, 2011). We speculated that the elevated ER stress levels in mutant livers might appear. Three main pathways, the PERK, IRE1, and ATF6 pathways, are known to mediate the ER stress response (also called the unfolded protein response/UPR). Each of these pathways evokes specific signal transduction and cellular reactions. Activating each ER-stress response/UPR pathway induces expression of a set of genes to resolve the stress in the ER. Using quantitative real-time PCR (qPCR), we examined the expression of several UPR-related genes, such as *xbp1s, chop, grp78*, and *grp94* to reveal which pathway(s) is/are activated in mutant fish. Total RNA was isolated from three-week-old larvae and the mRNAs were reverse transcribed. The genotype of each larva was determined by melt curve analysis using real-time PCR (Supplementary Fig. S2). In the non-stressed condition, transcribed *xbp1* pre-mRNA remains untranslated in the cytoplasm. Upon stress induction, ER membrane-bound Ire1 protein is phosphorylated, stimulating the splicing of *xbp1* to form *xbp1-s* mRNA. Subsequently, Xbp1-s protein is translated and activates its target genes. *chop* is one of the targets of the Perk pathway. Once ER membrane-bound Perk protein is phosphorylated upon ER stress, Atf4 protein translation is initiated via elF2-alpha protein activation. Atf4 in turn activates *chop*, which is a transcription factor for several genes that mediate apoptosis. Atf6 is another ER-membrane-tethered protein. Upon ER stress, Atf6 moves to the Golgi apparatus where it undergoes cleavage and is released from the membrane. The soluble ATF6 fragment translocates to the nucleus to mediate target gene activation, including ER-resident chaperones such as BiP (also known as grp78), and grp94, though these are also targets of the Ire1 pathway (Hetz, 2012; Oslowski and Urano, 2011). We found that expression levels of *chop* and *xpb1s* did not change, while those of *BiP*/*grp78* and *grp94* were upregulated in mutants (Fig. 4). These results indicate specific activation of the Atf6 pathway. Such partial activation of the ER-stress response/UPR suggests a constant elevation in ER-stress in the mutant that alters lipid metabolism in the ER, leading to liver defects with lipidosis.

### Failure of sex maturation in mutant fish

Even though some mutant fish can survive for several months, they do not show clear sexual differentiation. Because they lack clear hallmarks of gender, such as sex-specific body shape and fin color, inspection of internal organs by dissection was required to determine their sexes. When female fish were dissected, small ovaries containing premature oocytes were found (Fig. 5A). They lacked the ability to lay eggs when mated with wild-type or mutant males. When mutant male fish were dissected, we found normal-looking testes, and sperm motility was unaffected. Nevertheless, no mutant males succeeded in producing offspring when mated with wild-type females. We have not clearly identified the cause for male infertility, but postulate that it could be due to lack of neurological maturation, or under-developed physique that prevents their acceptance by females.

In human patients with ZS, skeletal abnormality such as enlarged fontanelles and spontaneous joint calcifications was reported. We, thus, examined skeletal phenotypes of mutant fish. Cartilages were stained by Alcian blue and bones were stained with alizarin red S. We did not find any clear morphological differences of skeletons between mutant fish and wild-type siblings (Fig. 5B).

### Mutant fish show a locomotive defect

A failure in locomotion appears in the patients with ZS and in ZS model mice. Similar to this, the mutant fish, which survived for some weeks, showed behavioral abnormality (Fig. 5C, Supplementary movie S1 and movie S2). Zebrafish normally swim around the fish tank, but the mutant fish frequently lay on the bottom of the fish tank and often failed to keep their upright posture when swimming. The phenotype indicates possible neuronal and/or muscular defects, which are also suggested in human patients and model animals (Takashima and Shimozawa, 2020).

### Genome-wide gene expression analysis revealed mutant-specific gene expression changes

To explore the transcriptomic effect of peroxisomal dysfunction, we performed a genome-wide gene expression analysis. We designed a custom microarray exclusively includes zebrafish genes with Refseq ID (Table S2; see Materials and Methods for detail). Gene expression was compared by the microarray between 5-week old mutant fish and their wild-type homozygous siblings. The summary of the microarray analysis is shown in Fig. 6 and Table S3. After moderated *t*-test and Benjamini-Hochburg multiple testing correction with a false discovery rate (FDR) cutoff of 0.05 (Benjamini and Hochberg, 1995; Smyth, 2004), we found 9373 genes with upregulated (5683) or down-regulated (3724) expression in mutant fish compared with wild-type siblings (34 genes had up- and down-regulated transcriptional variants). Of these, 4380 genes displayed ≥ 2-fold up (2142 genes) or down (2243 genes) regulation (5 genes displayed up and down regulated transcript variants), and 670 genes displayed ≥ 5-fold expression change (62 upregulated and 608 downregulated, with no overlap) (Table S3). Compared to up-regulated genes, down-regulated genes showed greater degree of expression changes (Fig. 6D).

**Fig 6.**
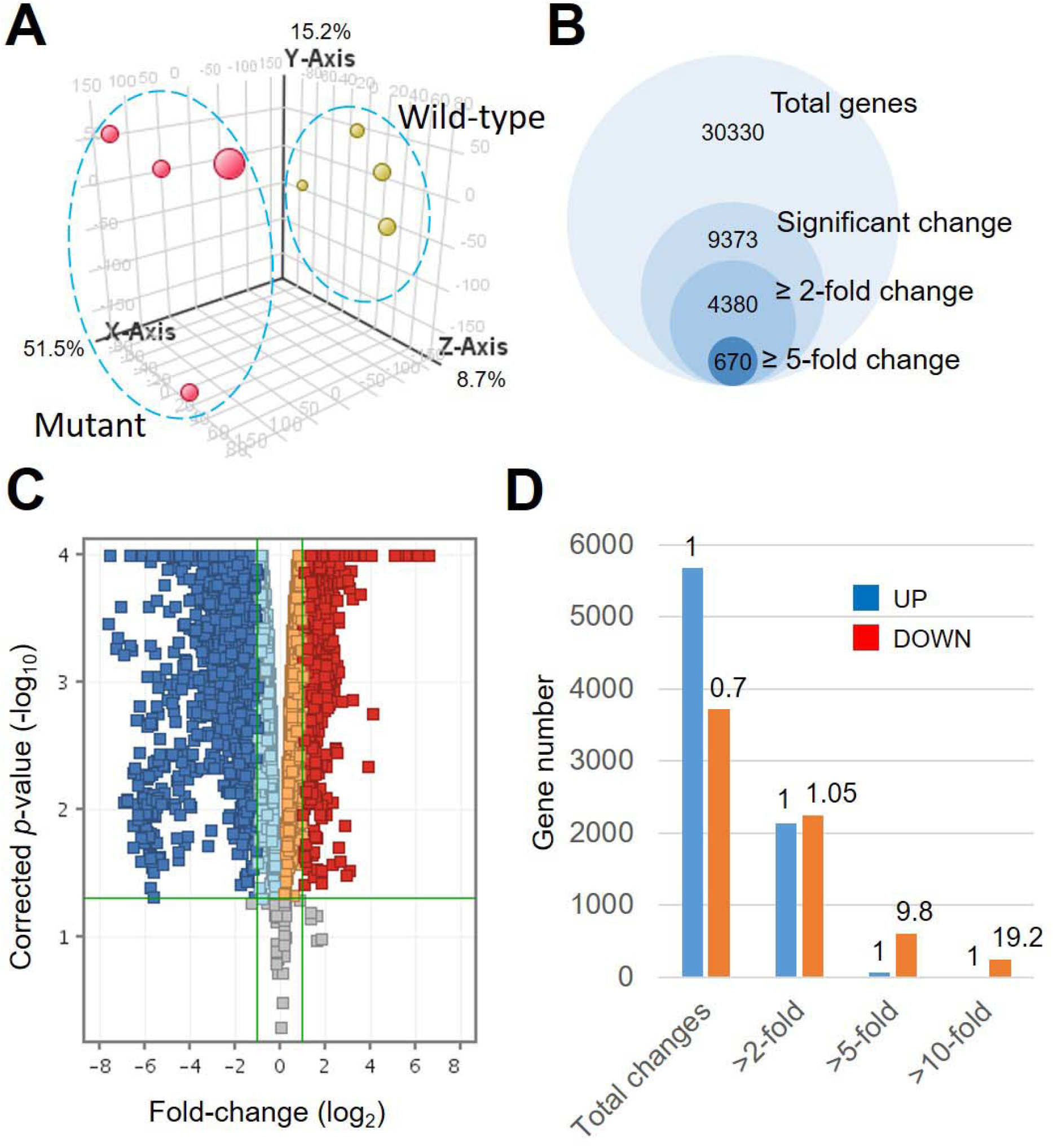
Overview of microarray analysis. (A) Principal component analysis of microarray data. The first three principal components of the data (represented by *x, y*, and *z* axes, respectively) were shown. The numbers beside each axis indicate the cumulative contribution rate. Mutant data were shown in red and wild-type data were in yellow. The two groups were clearly segregated each other mainly along the first component (*x* axis). (B) Venn diagram of genes detected by each analytical criterion. Numbers of total genes on the microarray, genes with significant expression change, genes with ≥ 2-fold expression change, and genes with ≥ 5-fold expression change were shown. (C) Volcano plot showing genes with significant expression changes with ≥ 2-fold upregulation (red) and downregulation (blue). (D) Bar graph showing the number of genes with significant expression change, ≥ 2-fold, ≥ 5-fold, and ≥ 10-fold expression changes, is shown. The numbers above the bars indicate the ratio of down-regulated genes to up-regulated genes in each class.

We performed functional annotation analysis of genes with altered expression using DAVID (see Materials and Methods), and found remarkable changes in specific biological features. Table 2 summarizes the results of GO term enrichment analysis using genes that showed ≥ 2-fold downregulation. All but one of the top 25 GO terms can be categorized into four categories, namely “gamete development”, “cellular chemotaxis”, “muscle contraction”, and “inflammatory responses”. The tissue specificity of the down-regulated genes was also analyzed, and testis, ovary, lens, gills, skin, and gut were found to express many of the genes (Table S4). We also performed KEGG pathway analysis using genes with ≥ 2-fold up- or down-regulation. Pathways related to “cell-cell interaction”, “lipid and fatty acid metabolism”, and “muscle contraction” are strongly affected in the mutants (Table S5 and Table S6). From these analyses, we identified biological events altered in the mutants. These include development of reproductive organs, muscle contraction, chemotaxis, inflammatory responses, lipid and fatty acid metabolism, and cell-cell interaction. We also identified primarily affected organs and tissues, which include ovary, testis, muscle, lens, gut, and skin.

**Table 2.** Gene Ontology (GO) analysis with ≥ 2-fold down-regulated genes.

| Category <sup>(1)</sup> | GO term | Count | Fold enrichment | *p*-value | FDR |
| --- | --- | --- | --- | --- | --- |
| GD | Germ cell development | 22 | 4.56 | 3.5E-09 | 6.9E-06 |
| GD | Female gamete generation | 18 | 5.51 | 4.8E-09 | 4.8E-06 |
| GD | Oogenesis | 17 | 5.60 | 1.1E-08 | 7.2E-06 |
| CC | Cell chemotaxis | 27 | 3.62 | 1.1E-08 | 5.4E-06 |
| - | Proteolysis | 116 | 1.62 | 1.3E-07 | 5.3E-05 |
| CC | Lymphocyte migration | 13 | 6.19 | 2.6E-07 | 8.8E-05 |
| CC | Leukocyte migration | 22 | 3.63 | 3.1E-07 | 8.8E-05 |
| MC | Heart contraction | 26 | 3.15 | 3.9E-07 | 9.7E-05 |
| GD | Gamete generation | 26 | 3.12 | 4.7E-07 | 1.0E-04 |
| CC | Leukocyte chemotaxis | 19 | 4.00 | 4.9E-07 | 9.7E-05 |
| IR | Response to interferon-gamma | 12 | 6.43 | 5.7E-07 | 1.0E-04 |
| CC | Neutrophil chemotaxis | 15 | 4.94 | 6.4E-07 | 1.1E-04 |
| MC | Striated muscle contraction | 15 | 4.82 | 9.1E-07 | 1.4E-04 |
| GD | Positive regulation of acrosome reaction | 9 | 8.90 | 1.2E-06 | 1.7E-04 |
| GD | Egg coat formation | 9 | 8.90 | 1.2E-06 | 1.7E-04 |
| GD | Binding of sperm to zona pellucida | 9 | 8.90 | 1.2E-06 | 1.7E-04 |
| GD | Sperm-egg recognition | 9 | 8.90 | 1.2E-06 | 1.7E-04 |
| CC | Granulocyte chemotaxis | 15 | 4.48 | 2.5E-06 | 3.3E-04 |
| IR | Cellular response to interferon-gamma | 11 | 6.15 | 3.4E-06 | 4.3E-04 |
| IR | Cytokine-mediated signaling pathway | 24 | 2.86 | 7.2E-06 | 8.4E-04 |
| CC | Neutrophil migration | 15 | 4.10 | 8.0E-06 | 8.9E-04 |
| IR | Cellular response to cytokine stimulus | 24 | 2.81 | 9.9E-06 | 1.0E-03 |
| IR | Response to tumor necrosis factor | 10 | 5.59 | 3.0E-05 | 3.0E-03 |
| MC | Skeletal muscle contraction | 10 | 5.36 | 4.5E-05 | 4.3E-03 |
| IR | Response to interleukin-1 | 10 | 5.36 | 4.5E-05 | 4.3E-03 |
| GD | Acrosome reaction | 9 | 6.09 | 4.5E-05 | 4.1E-03 |

<sup>(1)</sup> GD: gamete development, CC: cellular chemotaxis, MC: muscle contraction, IR: inflammatory responses.

We also analyzed gene-level alterations in organs. The ≥ 2-fold down-regulated genes in the testis and ovary are listed in Table S7. Of these, 121 genes were ovary specific, 35 genes were testis specific, and 27 genes were expressed in both organs. Dysfunction of genes expressed in both organs could cause failure of sex differentiation in both male and female mutant fish. In KEGG pathway analysis, the “progesterone-mediated oocyte maturation” pathway was indicated. Altered expression of genes in this pathway might cause loss of sex differentiation as well (Table S8). In muscle, down-regulated genes include many *troponin* and *myosin* genes, which are required for muscle contraction (Table 3), indicating that muscle contraction is affected by altered expression of these genes. In the lens, down-regulated genes included many *crystallin* genes (Table 4). We thus examined all 62 *crystallin* genes present in the zebrafish genome in a heat map, and found that half of them (32/62) were down-regulated by more than 2-fold in the mutant fish (Fig. 7). As shown above, the details of genetic alterations in mutant fish were elucidated.

**Table 3.** Genes in muscle contraction with ≥ 2-fold down-regulation.

| **Entrez**<br>**GeneID** | **Gene**<br>**Symbol** | **Gene Name** | **Fold**<br>**Change** |
| --- | --- | --- | --- |
| **402809** | ***tnni2a.3*** | ***troponin I type 2a (skeletal, fast), tandem duplicate 3*** | **-23.61** |
| 373074 | *dnd1* | *DND microRNA-mediated repression inhibitor 1* | -8.36 |
| **492493** | ***tnni2a.1*** | ***troponin I type 2a (skeletal, fast), tandem duplicate 1*** | **-7.85** |
| 100002038 | *si:rp71-17i16.4* | *si:rp71-17i16.4* | -7.37 |
| 553296 | *hrc* | *histidine rich calcium binding protein* | -7.25 |
| 556258 | *tcap* | *titin-cap (telethonin)* | -6.62 |
| **449825** | ***tnni4a*** | ***troponin I4a*** | **-6.34** |
| **406437** | ***tpm4a*** | ***tropomyosin 4a*** | **-5.54** |
| **554168** | ***myh11a*** | ***myosin, heavy chain 11a, smooth muscle*** | **-4.71** |
| 30393 | *cryaba* | *crystallin, alpha B, a* | -4.28 |
| **550249** | ***tnni2b.1*** | ***troponin I type 2b (skeletal, fast), tandem duplicate 1*** | **-4.08** |
| **64671** | ***cmlc1*** | ***cardiac myosin light chain-1*** | **-4.01** |
| **394068** | ***tnni2a.2*** | ***troponin I type 2a (skeletal, fast), tandem duplicate 2*** | **-3.88** |
| 503591 | *fbxl22* | *F-box and leucine-rich repeat protein 22* | -3.48 |
| 407664 | *hbegfb* | *heparin-binding EGF-like growth factor b* | -3.34 |
| **751665** | ***tnni1c*** | ***troponin I, skeletal, slow c*** | **-3.20** |
| 195815 | *lmna* | *lamin A* | -3.11 |
| **436902** | ***tnni1d*** | ***troponin I, skeletal, slow d*** | **-2.77** |
| **30592** | ***myl7*** | ***myosin, light chain 7, regulatory*** | **-2.77** |
| **58071** | ***tnnt2a*** | ***troponin T type 2a (cardiac)*** | **-2.55** |
| **494070** | ***tnni1b*** | ***troponin I type 1b (skeletal, slow)*** | **-2.51** |
| **353247** | ***tnnc1a*** | ***troponin C type 1a (slow)*** | **-2.33** |
| 322509 | *acta2* | *actin, alpha 2, smooth muscle, aorta* | -2.26 |
| 30696 | *nkx2.5* | *NK2 homeobox 5* | -2.18 |
| 393510 | *slc2a12* | *solute carrier family 2 (facilitated glucose transporter), member 12* | -2.10 |
| 450005 | *csrp3* | *cysteine and glycine-rich protein 3 (cardiac LIM protein)* | -2.07 |
| 415107 | *bves* | *blood vessel epicardial substance* | -2.02 |

*troponin* and *myosin* genes are shown in bold.

**Table 4.**
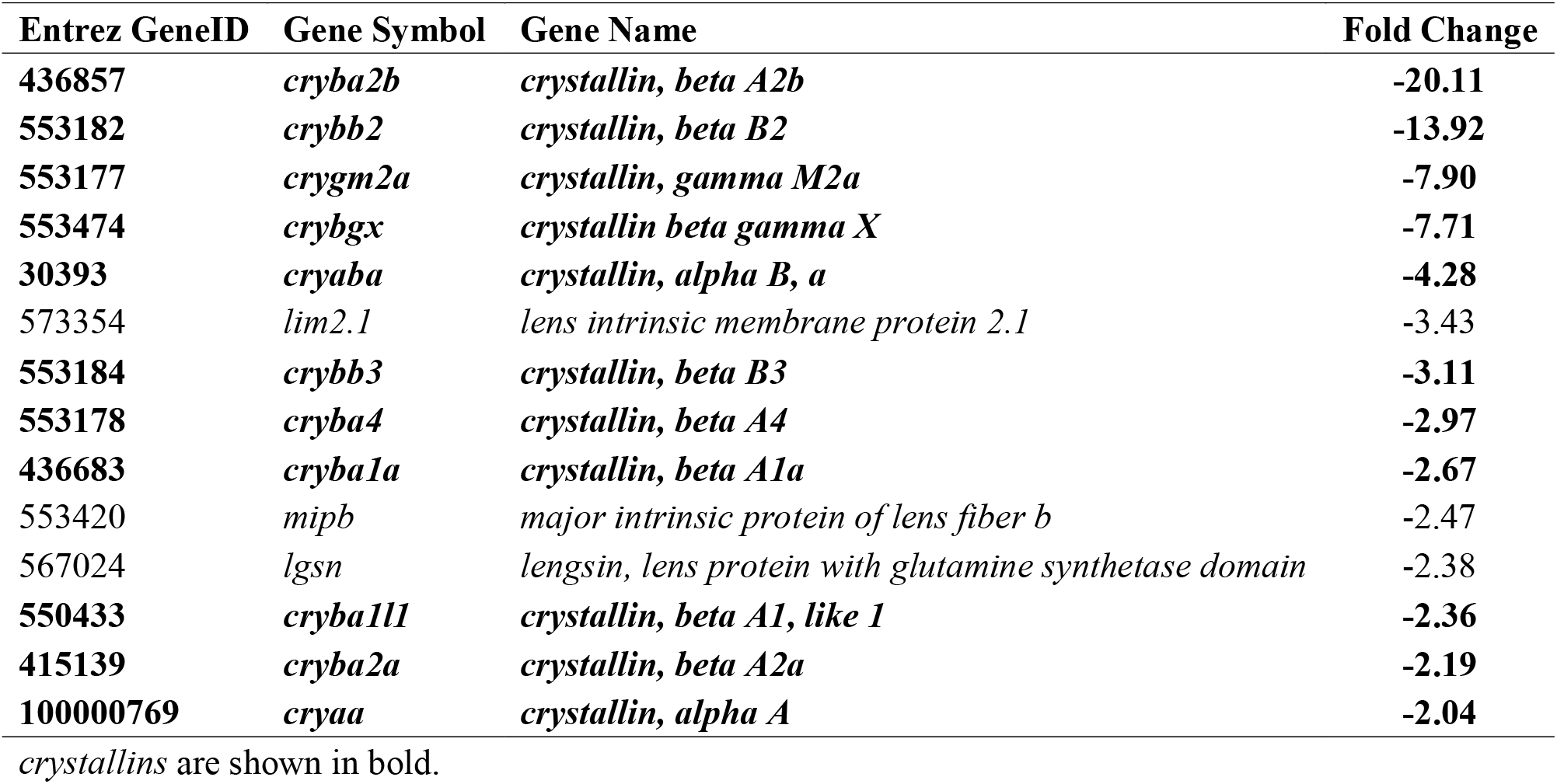
Lens expressed genes with ≥ 2-fold down-regulation.

**Fig 7.**
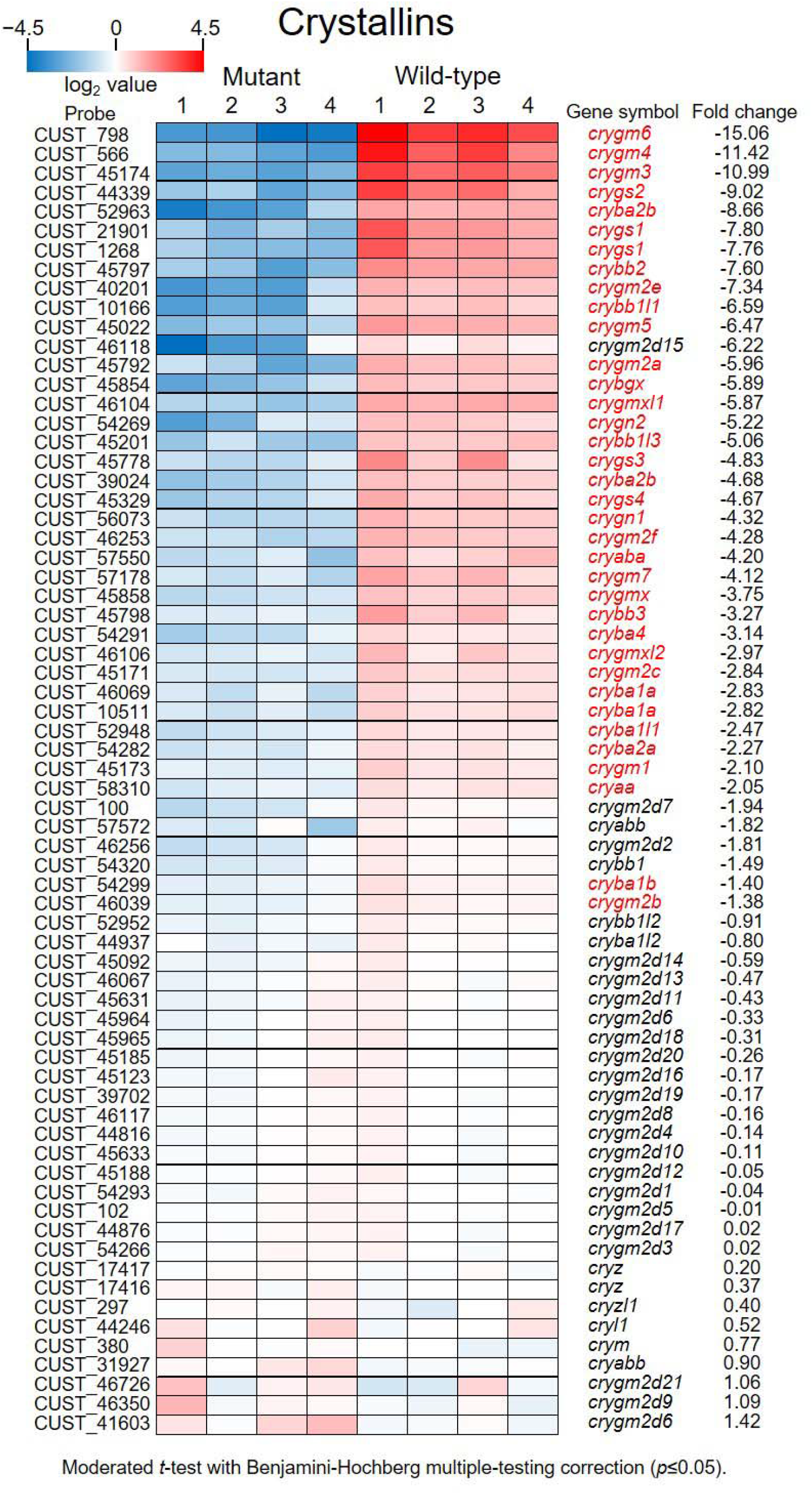
Crystallins are decreased in *pex2* mutant fish. A heat-map of *crystallin* genes. The expression levels of 62 *crystallin* genes are compared between *pex2* mutant fish and wild-type siblings by microarray. The number above the heat-map represents the array number (each array contains 3 individuals). Gene symbols in red represent genes with significant expression changes revealed by statistical analysis (see Materials and Methods for details). Negative fold change values indicate reduced gene expression in mutant fish.

We also analyzed selected groups of genes and gene families. In the zebrafish genome, 67 genes are listed in the KEGG pathway “fatty acid metabolism” (KEGG pathway ID: dre01212) and 18 genes within this family encode peroxisomal proteins, whose expression would be affected in ZS (Fig. 8). Notable genes in this pathway with altered expression are *acyl-CoA synthetase long-chain family members* (*acsl*) *1a, 1b, 4b*, and *5*, which catalyze the synthesis of CoA-conjugated FAs for elongation and degradation of FAs. While *acsl1a* and *acsl1b* were upregulated in the mutants, *acsl4b* and *acsl5* were downregulated, suggesting that they would affect FA type-dependent metabolic pathways. The remarkable non-peroxisomal gene family is *elovl7a* and *7b*, which mediate VLCFA elongation, and are both down-regulated; *ppt1* (*palmitoyl-protein thioesterase 1*) and three *ppt2a* (*palmitoyl-protein thioesterase 2a, tandem duplicates 1, 2, and 4*) genes, which are required for removal of thioester-linked FAs from FA-conjugated proteins (Fig. 8).

**Fig 8.**
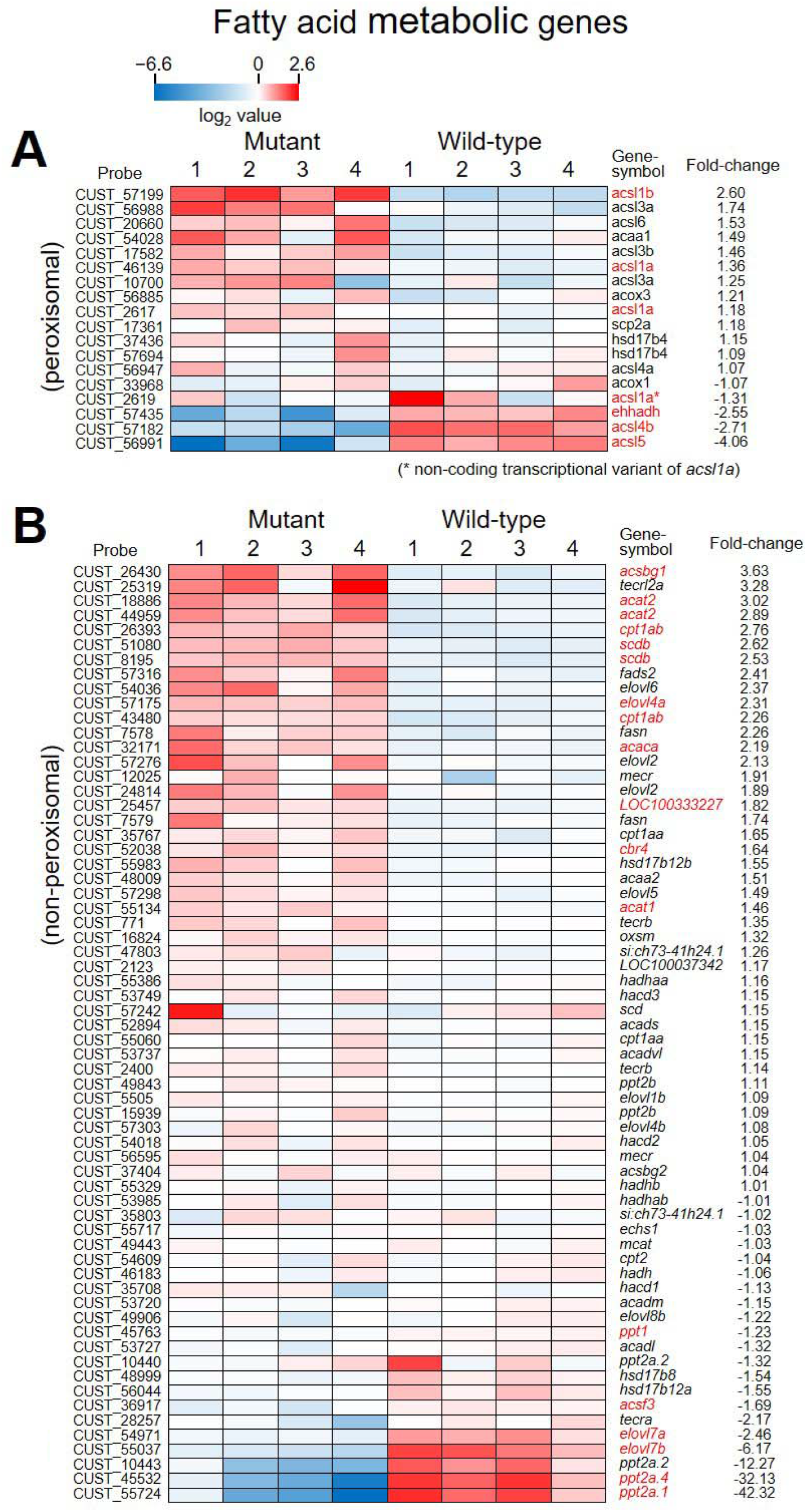
Expression of fatty acid metabolic pathway genes. (A, B) Heat-maps showing expression of genes related to fatty acid metabolism. Peroxisomal genes are shown in (A), and non-peroxisomal genes are shown in (B). Gene symbols in red represent genes with significant expression changes revealed by statistical analysis.

Parvalbumins (Pvalbs) are low-molecular weight albumins with calcium binding activity that are required for development and physiological activities of the central nervous system and musculature. In patients with ZS and in ZS model mice, failures in muscle and Purkinje cells, both express *Pvalbs*, are often reported (Faust et al., 2001; Krysko et al., 2007; Wanders, 2013). In the zebrafish genome, there are nine *pvalb* genes from *pvalb1* to *pvalb9. pvalb1, pvalb2, pvalb3, pvalb4*, and *pvalb7* are expressed in somites/myotomes (Boyle Anderson and Ho, 2018; Thisse and Thisse, 2004; Thisse et al., 2001), and *pvalb6* and *pvalb7* are expressed in Purkinje cells during embryonic development and in adult fish (Bae et al., 2009; Takeuchi et al., 2017). *pvalb3, pvalb4, pvalb5, pvalb7, and pvalb9* were found to be significantly downregulated in the mutant fish (Fig. 9), suggesting possible defects in muscle function and Purkinje cell development in mutant fish.

**Fig 9.**
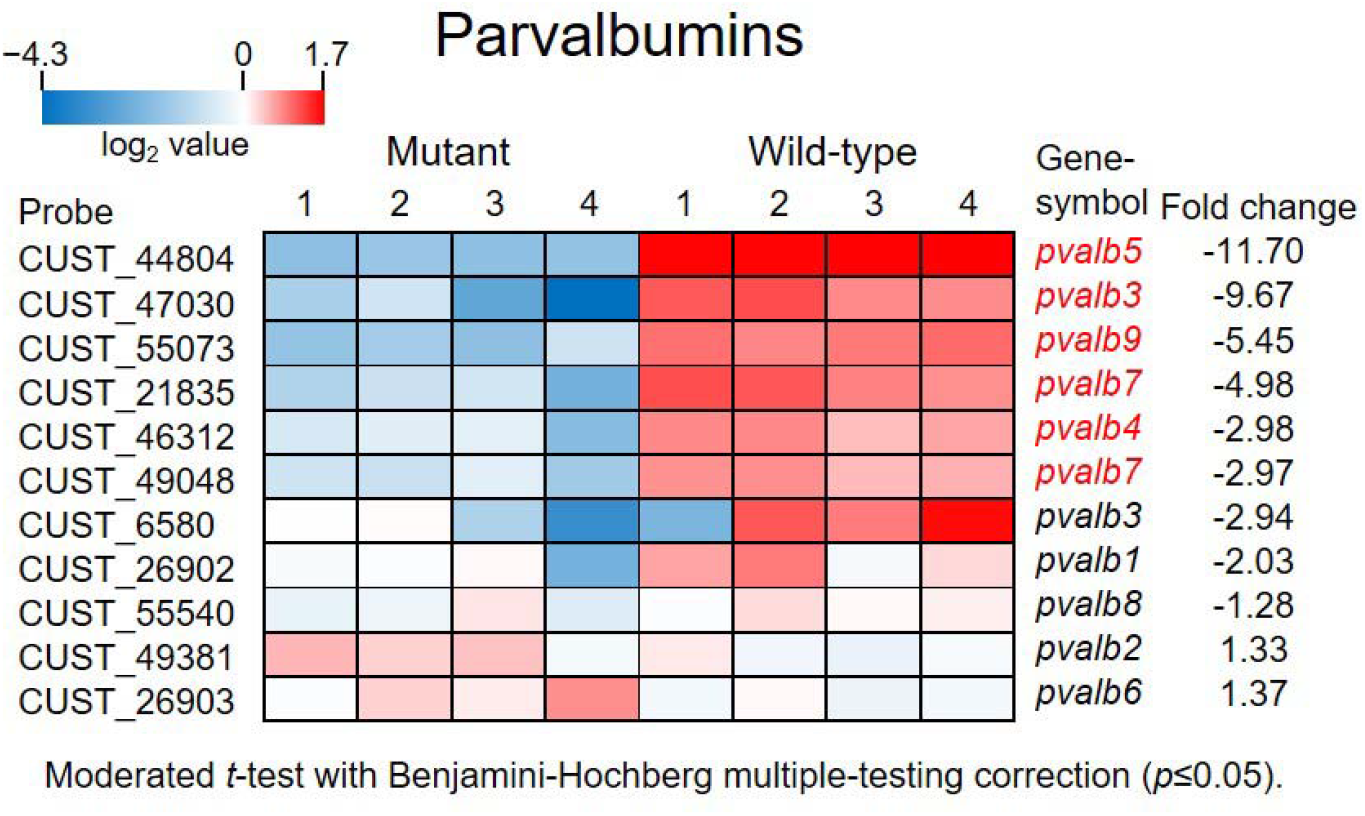
Comparison of Parvalbumin expression. A heat-map of nine *parvalbumin* genes. *parvalbumin* gene expression levels are compared by microarray analysis. Gene symbols in red represent genes with significant expression changes revealed by statistical analysis. Negative fold change values indicate decreased gene expression in mutant fish.

Due to limited knowledge of miRNAs in ZS, we analyzed whether miRNA expression was affected in mutant fish. Our custom microarray includes 283 miRNA probes, although they actually target the poly(A) tail-retaining pri-miRNAs. We found that 63 miRNAs were upregulated and 12 miRNAs were downregulated in the mutant fish. Of these, 21 and 11 displayed more than 2-fold expression changes, respectively (Supplementary Fig. S3). Changes in the miRNA profile imply widespread post-transcriptional alterations in gene functions via miRNA-mediated translational attenuation in the mutant fish, although the miRNA targets remain largely unknown.

## DISCUSSION

ZS is a prototypical peroxisomal disorder characterized by total loss of functional peroxisomes and the most severe symptoms (Klouwer et al., 2016; Shimozawa, 2020a). We established the first genetically modified ZS model fish with targeted gene knock out, and showed that the ZS model fish lose peroxisomal biogenesis and develop similar symptoms as human patients, including locomotive defect, eating disability, liver abnormality, and early death. The mutant fish also showed increased levels of tissue VLCFAs and branched fatty acids, and a decrease in ether phospholipids, mimicking metabolic changes seen in human patients. *Pex* genes have been mutated in other model organisms, such as mice and fruit flies, to generate ZS model animals. In mice, all twelve disease-causing *Pex* genes are conserved, and five of them (*Pex1, Pex2, Pex5, Pex10*, and *Pex13*) have been targeted for KO (Baes and Van Veldhoven, 2012; Takashima and Shimozawa, 2020). Model mice have recapitulated human symptoms such as hypotonia, sucking disability, liver abnormality, neural migration defect in the cerebrum, abnormal Purkinje cells, and neonatal mortality. Most of the disease-causing *pex* genes, except for *pex26*, exist in the fruit fly genome, and their functions in peroxisomal biogenesis have been confirmed in cell line studies (Mast et al., 2011). To date, seven *pex* genes (*pex1, pex2, pex3, pex10, pex12, pex16*, and *pex19*) have been inactivated in the fly, and some of these mutant flies have recapitulated human symptoms (Takashima and Shimozawa, 2020). We discuss our findings relative to those found in other ZS model animals in the following sections.

In the livers of our ZS model fish, lipid droplets accumulate in hepatocytes. The lipid droplets enlarge as the mutant fish grow, and the shape of the liver becomes deformed. Hepatic steatosis is also common in many ZS mouse models (Baes and Van Veldhoven, 2016; Takashima and Shimozawa, 2020). The *Acox1*-deficient mouse model, which is defective in VLCFA β-oxidation allowing VLCFAs to accumulate in the body, develops hepatic steatosis as well, suggesting that VLCFA accumulation may trigger this symptom (Fan et al., 1996; Sheridan et al., 2011). In our study, along with VLCFA accumulation, elevated ER stress was observed in mutant fish. Because ER stress is considered to be a strong inducer of nonalcoholic steatohepatitis (NASH) (Han and Kaufman, 2016; Lebeaupin et al., 2018; Malhi and Kaufman, 2011), the accumulation of VLCFAs might induce liver steatosis by activating ER stress. Similar observations have been reported in *pex2* KO mice (Kovacs et al., 2009).

Hypotonia is a very remarkable symptom in ZS patients. Infants with ZS show poor muscle tone. The same phenotype has been observed in model animals including mice, flies, and our ZS model fish. In mouse models with systemic *Pex* gene KO (i.e., the *Pex* gene is inactivated throughout the entire body), neonatal mice were severely hypotonic and growth retarded, resulting in neonatal death within days of birth (Baes et al., 1997; Faust and Hatten, 1997; Li et al., 2002; Maxwell et al., 2003). Flies with mutations in *pex1, pex3*, and *pex19* are hypotonic and tend to die early (Bülow et al., 2018; Faust et al., 2014; Mast et al., 2011; Nakayama et al., 2011). The cause of death is, in part, due to inability to eat, preventing flies from obtaining sufficient energy from their diet. ZS model fish display similar defects in eating ability and locomotion. Hypotonia can be caused by neuronal defects and/or muscular dysfunction. In gene expression analysis, we found altered expression of genes related to muscle contraction, especially strong down-regulation of *troponin* and *parvalbumin* genes, both of which encode distinct types of calcium-binding proteins or related proteins. Our gene expression data suggest that downregulation of *troponin* and *parvalbumin* genes might be the cause of muscular dysfunction. Their expression levels should thus be addressed in other animal models to understand the molecular nature of hypotonia in ZS patients and model animals.

We found that ZS model fish failed to mature sexually. None of the mutant fish, male or female, were fertile, even when mated with wild-type fish. In mutant females, oogenesis was clearly inhibited. Their ovaries were full of under-developed oocytes. By contrast, male fish have mature testes with normal-looking, motile sperm. We think, therefore, that the reason for infertility differs between males and females. It is possible that mutant males fail to develop sexually mature behaviors (i.e. neural circuits required for mating behavior may fail to develop). Flies with *pex2, pex10, pex12*, or *pex16* mutations were all viable with slight growth delays (Chen et al., 2010; Nakayama et al., 2011). They were infertile and showed clear defects in spermatogenesis. Because ZS model mice die before adulthood, their fertility and possible defects in gametogenesis have not been described. In the microarray analysis, we found clear gametogenesis pathway inhibition related to both sexes. Sex maturation requires sex hormones such as estrogens. We found that genes in the progesterone-induced oocyte maturation pathway were downregulated in ZS model fish, suggesting that reduction in sex hormone levels could cause infertility by affecting the gametogenesis pathway and/or the maturation of sex-related neural circuits.

In contrast to the mutant flies with non-lethal phenotypes, *pex1, pex3*, and *pex19* mutations were highly lethal in flies, producing neural defects in the CNS and PNS, reminiscent of vertebrate ZS models and human patients (Bülow et al., 2018; Faust et al., 2014; Mast et al., 2011; Nakayama et al., 2011). In most fly mutants, accumulation of VLCFAs has been reported. Interestingly, administration of medium-chain fatty acids (C12:0 and C14:0) ameliorate the lethality of *pex19* mutant flies, possibly by reducing mitotoxic free FAs (Sellin et al., 2018). Though it is unclear whether the cause of the lethality is the same in flies and vertebrates, the positive effect of MCFAs should be addressed in vertebrate models.

The relationship between peroxisomal metabolites such as VLCFAs, phytanic acid, or plasmalogens, and disease pathology remains unclear. In mice, *Pex5* function was attenuated in an organ-specific manner, which revealed that loss of peroxisomal function in the brain is not the primary cause of neural migration defects in the cerebrum or failure of Purkinje cell differentiation when liver peroxisomes are functional and blood metabolite levels are normal (Janssen et al., 2003; Krysko et al., 2007). This suggest that circulating metabolites are possible pathogenic agents. To date, the exact metabolites causing the symptoms have not been identified. We previously analyzed the FA profile in fibroblasts from ZS patients, and detected over 50 FA species in control fibroblasts and more than 100 species in patient fibroblasts (Takashima et al., 2017). In patient fibroblasts, uncommon VLCFA species were found. Saturated fatty acids reached chain lengths of 32 (C32:0), and unsaturated FAs reached C44 with many double bonds (for example, C44:13), which we call ultra-very long chain poly unsaturated fatty acids (ultra-VLC-PUFA). In ZS model fish, we also found organ-specific accumulation of ultra-VLC-PUFAs, such as C36:8, C36:9, C38:8, in the brain and eyes, but less in the liver. By contrast, the liver prominently displayed accumulation of shorter VLCFAs. This organ-specific accumulation of distinct FA species might be caused by organ-specific expression of FA metabolic genes, and resulting in organ-specific symptoms.

At the molecular level, we found remarkable gene expression changes that might be related to distinct symptoms in human patients. In mutant fish, reduced expression of many *crystallin* genes was revealed. Cataracts often develop in ZS patients by unknown mechanisms. Cataracts also appear in patients with RCDP type 1 through type 5, where limited peroxisomal metabolites such as plasmalogens and/or phytanic acid are affected, while VLCFAs are unaffected (Klouwer et al., 2016; Shimozawa, 2020b; Steinberg et al., 2006). In RCDP model mice, mutations in *Pex7* (RCDP type 1), *Gnpat* (RCDP type 2), or *Agps* (RCDP type 3) genes all result in development of cataracts (Braverman et al., 2010; Brites et al., 2003; Liegel et al., 2011; Liegel et al., 2014; Rodemer et al., 2003). Although expression of *crystallin* genes has not been characterized in ZS and RCDP patients and model mice, reduced expression of these important lens proteins might underlie cataracts in ZS and RCDP patients.

In this paper, we described phenotypes and gene expression changes in ZS model fish. The ZS model fish successfully recapitulated important phenotypes in human ZS patients, and altered levels of peroxisomal metabolites such as VLCFAs, branched fatty acids, and ether phospholipids. These findings indicate that peroxisomes share common functions among mammals and teleosts, and the pathological mechanisms are shared among vertebrate species. Because of high genetic tractability and accessibility of developing embryos and larvae, zebrafish offer an ideal ZS model in addition to mice and fruit flies to elucidate molecular details of pathology, and to establish and examine possible treatments for this devastating human disease.

## MATERIALS AND METHODS

### Zebrafish maintenance

A wild-type zebrafish strain (Riken WT, RW) was obtained from the National BioResource Project, Zebrafish (Riken, Saitama, Japan) and was used as the background strain for *pex2* mutant fish. Fish are maintained via standard methods in fish tanks at ≈ 28°C. A 14h light, 10h dark cycle was applied. Fish were fed brine shrimp and powdered meal multiple times daily. To increase viability, zebrafish larvae were fed *Paramecium tetraurelia* and *Paramecium sp*. (obtained from the National BioResource Project, Paramecium; Yamaguchi University, Yamaguchi, Japan) raised in barley grass extract (made from young barley grass powder; Yamamoto Kanpoh Pharmaceutical, Aichi, Japan), supplemented with *Klebsiella pneumoniae* (a gift from Dr. Hideo Dohra, Shizuoka University).

### TALEN-mediated genome editing

TAL effector nucleases (TALENs) were designed on ZiFiT (http://zifit.partners.org/ZiFiT/) and constructed using the Joung Lab REAL Assembly TALEN kit (a gift from Keith Joung; Addgene, Cambridge, MA, #1000000017), according to the provided protocol (Sander et al., 2011). The composition of the TALENs and their target sites in the zebrafish *pex2* gene are shown in Supplementary Fig. S4). TALEN mRNAs were synthesized using mMESSAGE mMACHINE T7 ULTRA Transcription Kit (Thermo Fisher Scientific, # AM1345). TALEN mRNA injection was performed using an air pressure injector (BIA-1, BEX, Tokyo, Japan) with a glass needle (GC100F-10; Harvard Apparatus, Cambridge, MA, USA).

### Genotyping

For genotyping of adult fish, tail-fins were clipped and dissolved in 50 mM NaOH at 95°C for 20 min. After cooling to room temperature, 0.1 volume of 1M Tris-HCl (pH 8.0) was added for neutralization. The resulting solution was used for PCR analysis. A pair of primers designed to amplify the wild-type *pex2* allele (wild-type primer set) or a pair of primers for mutant *pex2* allele (mutant primer set) were independently applied to PCR mixture to determine whether the fish had only wild-type or mutant alleles, or one of each (heterozygous). To ensure that the PCR was working in the case with primer pair unmatched to the template DNA (such as the mutant primer pair used with wild-type DNA), an upstream forward primer was also added to the PCR mixture to amplify a longer PCR product from both types of allele. Larvae used for genotyping were killed by instant chilling on ice or in a freezer, then posterior halves were dissolved with 25 mM or 50 mM NaOH. Anterior halves were used for immunostaining or fatty acid analysis.

### Survival rate assay

Genomic DNA was extracted from whole embryos, larval tissue, or fin clips from adult fish. The genotype of each individual was examined using the above method. Because dead embryos and larvae can be degraded and lost in petri dishes or the fish tank, the presumptive total number of mutant fish produced was estimated from the numbers of surviving non-mutant siblings. This number was then compared with the actual number of identified mutant fish to determine the survival rate.

### Immunohistochemistry

Livers were fixed with HistoChoice MB Tissue Fixative (VWR, Radnor, PA, USA, #H120) for 4 h at room temperature (RT), or overnight at 4°C. After washing out the fixative with PBT (PBS containing 0.02% Triton-X100), samples were blocked with 10% normal goat serum (NGS) diluted in PBT, then incubated with primary antibody overnight at 4°C. After washing with PBT, samples were incubated with a fluorophore-conjugated secondary antibody for 2 h at RT, or overnight at 4°C. After washing, samples were stained with DAPI (1μg/mL; diluted with PBT) for 30 min at RT, washed again, and mounted in Vectashield mounting medium (Vector Laboratories, Burlingame, CA, #H-1000). Specimens were examined and imaged using a confocal laser-scanning microscope (LSM710; Carl-Zeiss, Oberkochen, Germany). Images were processed with ImageJ (National Institute of Health, USA), Photoshop SE (Adobe, San Jose, CA, USA), and PowerPoint (Microsoft, Redmond, WA, USA). The following antibodies were used in this study (working dilution and other information in parentheses): rabbit anti-human catalase (1:500; Athens Research & Technology, Athens GA, #01-05-030000), rabbit anti-PMP70 (1:500; a gift from Dr. Tsuneo Imanaka, Hiroshima International University, and Dr. Masashi Morita, Toyama University, Japan), goat anti-rabbit IgG Alexa 488 (1:500; Thermo Fisher Scientific, Waltham, MA, USA, #A-11008), and goat anti-rabbit IgG Alexa 546 (1:500; Thermo Fisher Scientific, #A-11035).

### Oil Red O staining

Larvae were fixed with 4% formaldehyde/PBS overnight at 4°C. After washing with PBS-tw (PBS containing 0.1% Tween20) three times for 5 min each, samples were stained with 0.3% Oil Red O (ORO) solution (Sigma-Aldrich, #O1391) diluted in water for 15 min. After washing with PBS-tw, excess staining was removed by treating samples with 60% isopropanol twice for 5 min each. Samples were re-fixed with 4% formaldehyde/PBS for 10 min, washed with PBS-tw, and mounted in 50% glycerol. For counter staining, samples were incubated with DAPI (1μg/mL, diluted with PBS-tw) to stain nuclei, then mounted in Vectashield.

### Skeletal Preparations

Bones and cartilages were stained with alizarin red S (Sigma-Aldrich, #A5533) and Alcian blue (Muto pure chemicals, Tokyo, Japan, #40851), respectively, according to the protocol previously described (Takashima et al., 2007).

### Quantitative real-time PCR (qPCR)

Primers used in this study were designed using the Primer designing tool (https://www.ncbi.nlm.nih.gov/tools/primer-blast/) and listed in Table S1. Total RNA was isolated from whole mount larvae or dissected tissue using the Maxwell RSC simplyRNA Tissue kit (Promega Corp., Fitchburg, Wisconsin, USA; cat. #AS1340). First strand cDNA was synthesized using ReverTra Ace qPCR RT Master Mix (TOYOBO, Osaka, Japan, #FSQ-201) or ReverTraAce qPCR RT Master Mix with gDNA Remover (TOYOBO, #FSQ-301). Quantitative real-time PCR (qPCR) was performed using THUNDERBIRD SYBR qPCR Mix (TOYOBO, #QPS-201) with StepOnePlus instrument (Thermo Fisher Scientific) using the relative standard curve method. When necessary, sample genotypes were identified by analyzing melting curves of the PCR products amplified with primers designed for the *pex2* transcript (z*Pex2*-qFwd and zPex2-qRev primers), as shown in Supplementary Fig. S2. Data were statistically analyzed and graphs were prepared using EZR software (Kanda, 2013).

### Microarray analysis

Microarray analysis was performed by the Division of Genomics Research, Life Science Research Center, Gifu University. Total RNA was isolated from whole bodies of 5-week old mutant fish and their wild-type siblings using Maxwell RSC simplyRNA Tissue Kit. The quality of total RNA was evaluated using 2100 Bioanalyzer system (Agilent Technologies, Santa Clara, California, USA) and total RNA samples with RIN of 7 or higher were applied to the next step. Three RNA samples were mixed in equal amounts to make each RNA pool. Four pools from mutant fish and four pools from wild-type siblings were used for microarray analysis (n = 4 each from 12 individuals, respectively). Cy3-labelled probes were prepared from RNA using Low Input Quick Amp Labeling Kit 1-color (Agilent Technologies, #5190-2305) and hybridized with the custom microarrays (ID:085740, 8 x 60k arrays; Agilent Technologies) for 17h at 65°C. The microarray slide was washed and scanned with a microarray scanner (ArrayScan, Agilent Technologies) to obtain the fluorescent signals of the probes. Signals were processed for digitization using Feature Extraction software (Agilent Technologies) and analyzed with GeneSpring GX software (Agilent Technologies) for gene expression. Data analysis was performed as follows. Gene expression data were normalized by 75 percentile shift. Probes ranked in the lower 20 percentiles in all samples were filtered out. Probes flagged as “compromised” in all samples were filtered out. Probes showing a CV value of more than 50% in each condition (mutant or wild-type) were filtered out. A moderated *t*-test with the Benjamini-Hochberg method for multiple testing correction was used to reveal the significant differences in gene expression between the two conditions (Benjamini and Hochberg, 1995; Smyth, 2004). A corrected *p*-value cutoff (*i*.*e*., FDR) of 0.05 was applied. For further data interpretation, we used the Database for Annotation, Visualization and Integrated Discovery (DAVID; https://david.ncifcrf.gov/) and the Kyoto Encyclopedia of Genes and Genomes (KEGG; https://www.genome.jp/kegg/).

### Liquid chromatography mass spectrometry (LC-MS)

Quantitative analysis of peroxisomal metabolites was performed as follows. Tissue samples were homogenized in 400 μL of acetonitrile using ZircoPrep Mini beads (Nippon Gene, Tokyo, Japan; #FGZ50M). Fifty microliters of 5 N hydrochloride was added to each homogenate and incubated at 100°C in an oil bath for 1h. One hundred microliters of methanol, 400 μL of *tert*-BME, and 400 μL of water were added and mixed by vortexing for 1 min. After centrifugation at 190 × g for 5 min, the upper phase was collected, added to 800 μL of water, mixed by vortexing for 1 min, and centrifuged again. The upper phase was collected in a sample vial (GL sciences, Tokyo, Japan; #1030-51023) with a polytetrafluorethylene (PTFE) -coated cap (Membrane Solutions, Auburn, WA, USA; #LBSV412CSSLBS), and the organic phase was evaporated under a stream of nitrogen gas. Samples were dissolved in acetone and applied to LC-MS. We used a Waters Acquity ultra-performance liquid chromatography (UPLC) system equipped with Xevo QTof mass spectrometer. Waters Acquity UPLC BEH C8 column (2.1 × 50 mm, particle size 1.7 µm, pore size 130 Å, #186002877) with BEH C8 VanGuard Pre-Column (2.1 × 5 mm, particle size 1.7 µm, pore size 130 Å, #186003978) was used. The aqueous mobile phase A was purified water containing 1 mM (0.0077% w/v) ammonium acetate and 5.78 mM (0.01% w/v) ammonia. The organic mobile phase B was 100% acetonitrile. The linear gradient was as follows (indicated as the concentration of mobile phase B, % B): initial condition 20% B; 20% B to 95% B from 0min to 50min; held at 95% B until 52.5min, 20% B from 52.5min to 55min. Flow rate was 0.4 mL/min, and column temperature was 40°C. Injection volume was 5 µL. Analytes were ionized by negative-ion electrospray ionization (negative ESI) and mass spectra were obtained by time-of-flight MS. The scan range was 100–1000 *m/z*. The MS settings were as follows: capillary voltage, 1.0 kV; sampling cone, 56 (arbitrary value); extraction cone, 4.0 (arbitrary value); source temperature, 125°C; desolvation temperature, 350°C; cone gas flow, 60 L/h; and desolvation gas flow, 1000 L/h. The obtained data were analyzed using Masslynx and Quanlynx software (Waters), and graphs and charts were produced using Excel (Microsoft). Significant increases or decreases in each FA species were calculated by Student’s *t*-test and Benjamini-Hochberg multiple testing correction using Excel.

## Acknowledgements

We thank Ms. Yoshiko Wakihara, Mayumi Sumi, Yuki Yokoyama, and the Life Science Research Center for performing the microarray analysis and DNA sequencing, Dr. Masahiro Fujishima and National Bio Resource Project, Paramecium for providing *paramecia*.

## Supplementary movie S1

The swimming behavior of wild-type fish.

## Supplementary movie S2

The swimming behavior of *pex2*^*-/-*^ mutant fish

